# Effect of oncolytic virotherapy on immune microenvironment in immune subtypes identified in gastric cancer and hepatocellular carcinoma

**DOI:** 10.1101/2023.01.17.524374

**Authors:** Ziyi Wang, Shuguang Peng, Xi Chen, Zhen Xie, Shao Li

**Author notes:** Corresponding authors: Shao Li,; Zhen Xie.

## Abstract

Tumor occurrence and progression are significantly influenced by immunity, and the immune infiltration and immune-related gene expression in solid tumors are closely correlated to the response of patients to immunotherapy. In this study, the level of tumor infiltrating immune cells in gastric cancer and hepatocellular carcinoma samples from the TCGA database were assessed using ssGSEA, and the tumor samples were divided into two subtypes (Imm_H and Imm_L) with different immune cell infiltration level. The differences in immune cell percentage and immune checkpoint gene expression between the two subtypes indicated that the Imm_H group had higher levels of immune infiltration, but also more infiltrated immunosuppressive cells and higher mRNA levels of immune checkpoint genes. Then the immune subtype-specific gene network was built and the main modules representing the genes and functions that differ between the two immune subtypes were identified. To explore the effect of oncolytic virus on tumor immune microenvironment, we constructed the previously developed synthetic adenovirus containing the synthetic sensory switch gene circuit, assessed the antitumor effect in mouse models, and measured the proportion of different cell types by single-cell RNA sequencing. The results showed that synthetic oncolytic virus inhibited tumor development and altered the proportion of infiltrating immune cells, suggesting that synthetic oncolytic virus may have different mechanism on the two immune subtypes.

**Graphical abstract:** 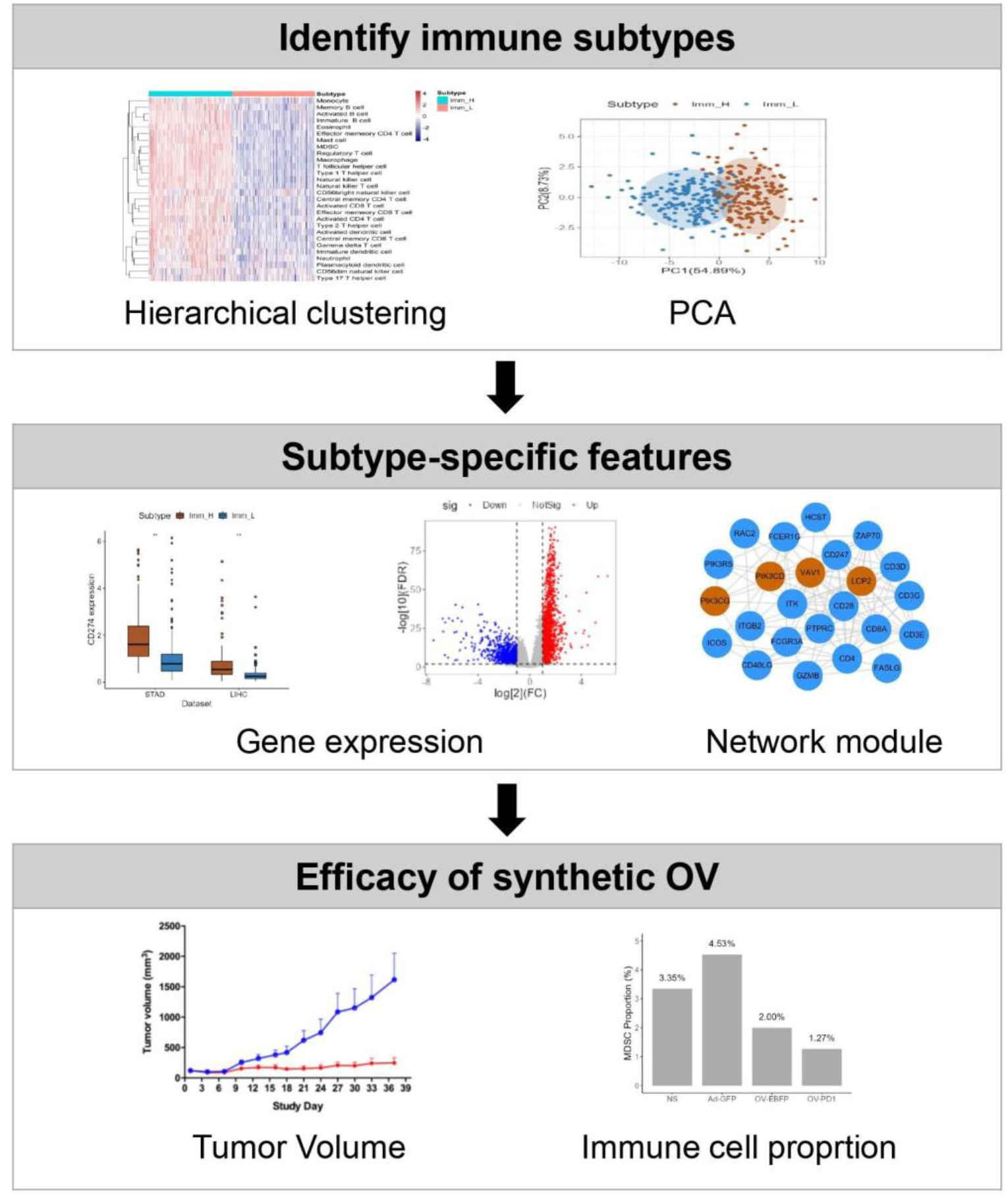

## 1. Introduction

Gastric cancer (GC) and hepatocellular carcinoma (HCC) are common cancers which account for a significant proportion of new cases and deaths from all cancers [1]. The development of tumor is a complex process in which multiple factors act synergistically [2]. Among them, the tumor microenvironment is an important factor that plays an important role in a variety of tumors [3–6]. Immune response in microenvironment is important in tumorigenesis and is related to tumor response to immunotherapy [7,8]. In the tumor microenvironment (TME), different kinds of immune cells as well as cytokines show anti-tumor or pro-tumor effects respectively. For example, natural killer (NK) cells have effective cytotoxic activity and inhibit tumor growth and metastasis. The presence of high density NK cells infiltration is related to good prognosis in various kinds of human solid tumors [9]. After T cell receptor-mediated signaling pathway is activated, regulatory T cells secrete inhibitory cytokines to promote T cell exhaustion and inhibit beneficial anti-tumor immunity [10]. As an innovative therapy, cancer immunotherapy has emerged as a focus of cancer therapy research. However, immunotherapy for GC and HCC still faces many challenges, such as the fact that it only succeeded in a limited number of patients and that it is difficult to predict the outcome.

Oncolytic virus (OV) has been recognized as an effective immunotherapy for cancers, which utilizes native or genetically modified viruses for selective replication in tumor cells [11]. OVs promote anti-tumor responses through two mechanisms that includes selective tumor lysis and induction of systemic anti-tumor immunity [12]. Additionally, immune modulators like granulocyte– macrophage colony-stimulating factor (GM-CSF) can be engineered to be expressed in OVs [13]. These immune modulators recruit circulating neutrophils, monocytes, and lymphocytes in a paracrine manner and enhance their functions in triggering anti-tumor immunity. Recombinant virus technology greatly improved the development OVs in tumor therapy, such as adenovirus, reovirus, herpes simplex virus, vaccinia virus, newcastle disease virus and coxsackie virus [11,14]. As of now, four OVs have been approved by different courtiers to cancer treatment, including Rigvir in Latvia, Oncorine (H101) in China, T-VEC (talimogene laherparepvec) in USA and Delytact (teserpaturev/G47Δ) in Japan [15–18].

To improve efficacy and tumor specificity of oncolytic virotherapy, we recently developed a platform to quickly construct synthetic adenovirus loading a synthetic sensory switch gene circuit [19]. The sensory switch gene circuit can sense and integrate multiple inputs i.e., tumor-specific promoters and microRNAs to regulate the viral replication and immune effectors release in the target cancer cells.

In this study, we downloaded and analyzed the gene expression profile of GC and HCC samples from the Cancer Genome Atlas (TCGA) and classified the tumor samples into two different subtypes according to their immune infiltration level. We analyzed the differences in immune characteristics between the two subtypes and identified subtype-specific genes and networks. Then we constructed oncolytic viruses loading synthetic sensory switch gene circuit, measured the inhibition of synthetic oncolytic virus on tumor growth and its effect on tumor microenvironment in mouse tumor models. The results showed that synthetic oncolytic virus has therapeutic effects on two tumor subtypes in different ways.

## 2. Material and methods

### 2.1 Data acquisition

The gene expression data (FPKM and read counts) and clinical information of GC patients and HCC patients were obtained from TCGA (https://cancergenome.nih.gov/). There were 375 gastric cancer samples and 32 normal samples in the TCGA-STAD dataset, and 371 hepatocellular carcinoma samples and 53 normal samples in the TCGA-LIHC dataset.

### 2.2 Differential expression analysis and enrichment analysis

Differentially expressed genes (DEGs) with q-value < 0.01 and | Log2 (Fold change) | > 1 between samples in different groups were identified by the EdgeR package [20]. According to the requirements, DEGs in tumor and paired normal samples, and DEGs in different immune infiltration subtypes were obtained. Gene ontology (GO) enrichment analysis was carried out using ClusterProfiler [21] to investigate the potential functions of DEGs. Gene set enrichment analysis (GSEA) was carried out using the GSEA R package [22], and KEGG pathways with q-value < 0.05 and | NES | > 1 were identified as significantly enriched pathways.

### 2.3 Grouping of cancer samples based on immune gene sets

Grouping of each cancer dataset was realized by single-sample gene-set enrichment analysis (ssGSEA). An immune-related gene set consisting of type-specific pan-cancer metagenes of 28 subpolulations of immune cells was collected from an earlier work [23]. For each tumor sample, the relative infiltration levels of each immune cell subgroups were quantified using the enrichment score in ssGSEA. Then, using hierarchical clustering, tumor samples from the TCGA-STAD and TCGA-LIHC were divided into two groups (Imm_H and Imm_L) according to the degree of immune infiltration.

### 2.4 Analysis of immune cell infiltration

Two independent methods were used in order to estimate immune cell infiltration in the Imm_H and Imm_L groups. Using the ESTIMATE algorithm [24], we calculated the stromal score, immune score, ESTIMATE score, and tumor purity of each tumor sample. Results showed the immune infiltration level and stromal content level in the tumor samples. In addition, we employed a deconvolution method CIBERRSORT [25] to identify fractions of immune subpopulations in tumor samples. The percentages of 22 different human immune cell subtypes were calculated after 1000 permutations.

### 2.5 Synthetic adenovirus construction

We constructed synthetic oncolytic viruses expressing different effectors using the previously developed assembly strategy [19]. We produced synthetic OVs encoding EBFP reporter (OV-EBFP), scFv against PD-1 (OV-PD1), human GM-CSF (SynOV1.1) and mouse GM-CSF (SynOV1.1m), respectively. Non-replicable adenovirus expressing GFP (Ad-GFP) was used as control.

### 2.6 Animal experiments

AGS tumors were initiated by SC implantation of 1 × 10^7^ AGS tumor cells into BALB/c nude mice. After tumor volumes reached approximately 120 mm^3^, two groups of animals (6 animals/group) received 3 IT injection of SynOV1.1 at the dose of 5.88 × 10^10^ VP or normal saline as negative control on Day 1, Day 8, and Day 15. mHepa1-6 tumors were initiated by SC implantation of 5 × 10^6^ mHepa1-6 tumor cells into the right flank of C57BL/6J mice. After tumor volumes reached approximately 100 mm^3^, mice were randomly divided into 4 groups (10 animals/group) and treated with saline as negative control, sorafenib as positive control at the dose of 60 mg/kg, SynOV1.1m at the doses of 5 × 10^8^ VP, and 5 × 10^10^ VP. The doses were administered intratumorally for SynOV1.1m on Day 1, Day 8, Day 15, and orally for sorafenib for 4 consecutive weeks (5 consecutive days on and 2 days off per week). Hepa1-6 tumors were initiated by SC implantation of 5 × 10^6^ Hepa1-6 tumor cells into the right flank of C57BL/6J mice. After tumor volumes reached approximately 200 mm^3^, mice were randomly divided into 4 groups and treated with saline as negative control, Ad-GFP, OV-EBFP, OV-PD1 at the doses of 5 × 10^9^ VP. Tumors were dissected for cell digestion, CD45 magnetic bead sorting and single-cell sequencing on the 10th day after treatment.

## 3. Results

### 3.1 Transcriptome analysis of paired tumor and paracancerous samples

To characterize the transcriptional profile in cancer and normal samples, we first obtained the transcriptome data of gastric cancer, hepatocellular carcinoma and corresponding paracancerous normal samples from TCGA database, and identified the DEGs between the paired tumors and paracancerous samples. 4656 and 2217 DEGs were identified from the TCGA-STAD and TCGA-LIHC datasets, respectively (Fig. 1A). GO enrichment results showed that DEGs from both datasets were significantly enriched in cell proliferation related GO terms and immune related GO terms, including DNA replication, nuclear chromosome segregation, antigen processing and presentation, leukocyte migration, and regulation of T cell activation (Fig. 1B-1C). Although only 14.9% of all DEGs are shared in gastric cancer and hepatocellular carcinoma (Fig. 1A), their enriched biological functions are similar and both reflected the excessive proliferation of cancer cells and the altered immunity in tumors. GSEA further revealed significant positive enrichment of cell cycle, p53 signaling pathway, antigen processing and presentation (NES>1), and significant negative enrichment of metabolic pathways, drug metabolism and cytochrome P450 (NES<−1) in both gastric cancer and hepatocellular carcinoma. In addition, KEGG pathways related to immune and inflammation such as Th17 cell differentiation, chemokine signaling pathway, and IL-17 signaling pathway were significantly positively enriched in gastric cancer samples. However, cytokine-cytokine receptor interaction was significantly negatively enriched in hepatocellular carcinoma samples (Fig. 1D-1E).

**Fig. 1.**
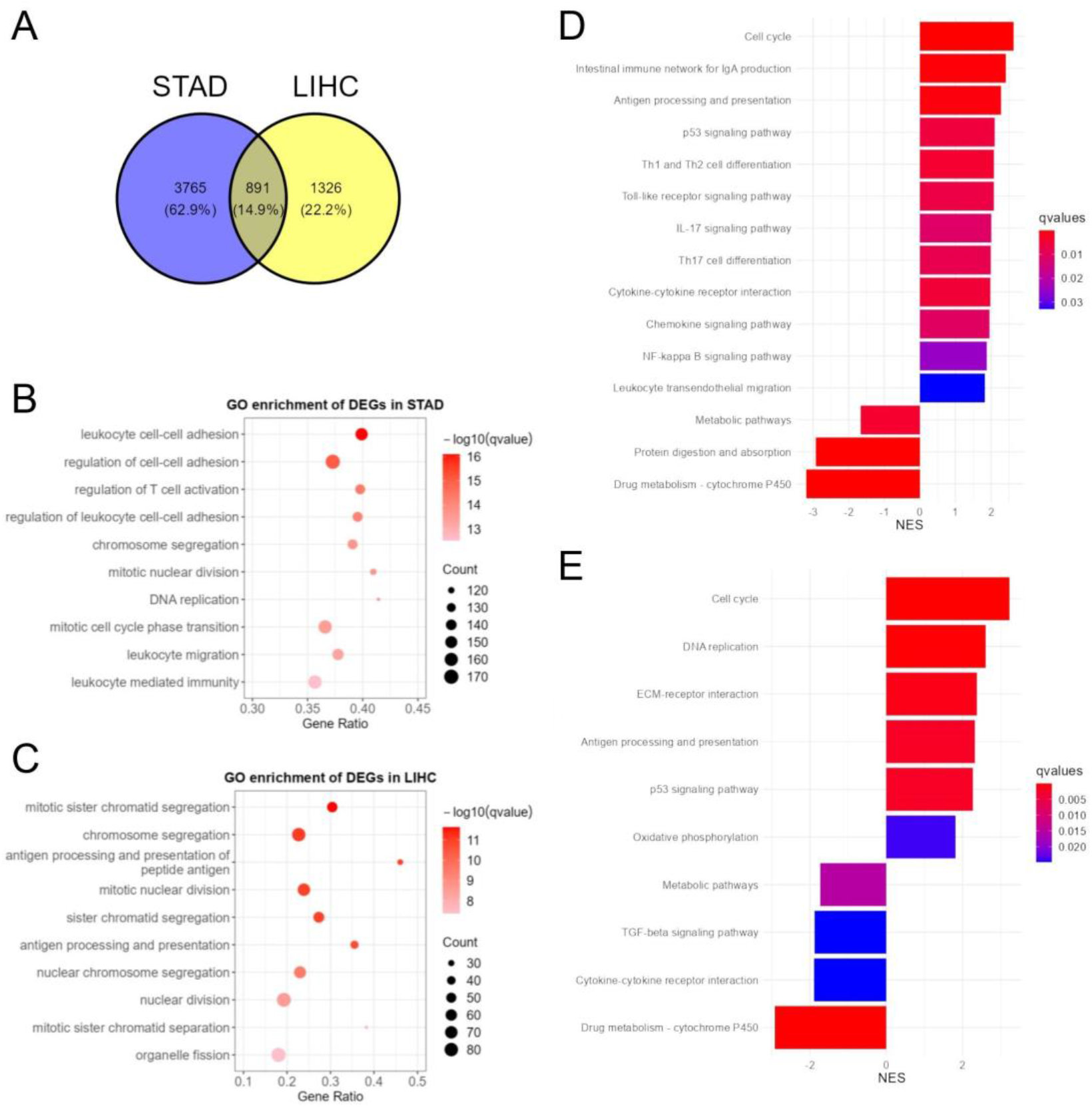
Differentially expression analysis and enrichment analysis between tumor samples and normal samples. (A) Venn diagram of DEGs in TCGA-STAD and TCGA-LIHC. Diagram shows that 891 DEGs were overlapped in TCGA-STAD and TCGA-LIHC. (B-C) GO term enrichment of DEGs in TCGA-STAD and TCGA-LIHC. (D-E) GSEA results in TCGA-STAD and TCGA-LIHC. Gene sets with |NES| > 1 and q-value < 0.05 were shown.

### 3.2 Identification and verification of immune subtypes in GC and HCC

Recent studies have demonstrated the importance of tumor-infiltrating immune cells in gastric cancer and hepatocellular carcinoma. Numerous studies have shown the complicated and varied functions that various immune cells play at different stages of different cancers, as well as their effects on tumor progression and prognosis. To investigate the immune microenvironment in GC and HCC, we downloaded transcriptional profiles of 375 stomach adenocarcinoma samples and 371 hepatocellular carcinoma samples from TCGA. The ssGSEA method was used to analyze 28 immune cell subtype-associated gene sets, and the ssGSEA score suggests the infiltration of different immune cell subsets in each sample. According to the ssGSEA scores, hierarchical clustering was performed on tumor samples from two TCGA datasets (STAD and LIHC). The tumor samples could be categorized into Imm_H group with higher immune cell infiltration and Imm_L group with lower immune cell infiltration (Fig. 2A-2B). Gastric cancer samples were classified into 188 samples in Imm_H group and 187 samples in Imm_L group, while hepatocellular carcinoma samples were classified into 120 samples in Imm_H group and 251 samples in Imm_L group. Principal component analysis based on the ssGSEA scores verified that the tumor samples could be divided into two subtypes (Fig. 2C-D). To further confirm the grouping results, the tumor purity, ESTIMATE score, immune score and stromal score of the two subtypes were calculated and compared. Violin plots showed that in comparison to the Imm_L group, the Imm_H group exhibited significantly higher immune score, stromal score and ESTIMATE score in both datasets (P<0.001). In contrast, the tumor purity in the Imm_H group was significantly lower than that in the Imm_L group (P<0.001) (Fig. 2D-2E). According to these findings, the tumor samples in the Imm_H group had more immune cells and stromal cells, and those in the Imm_L group contained more tumor cells.

**Fig. 2.**
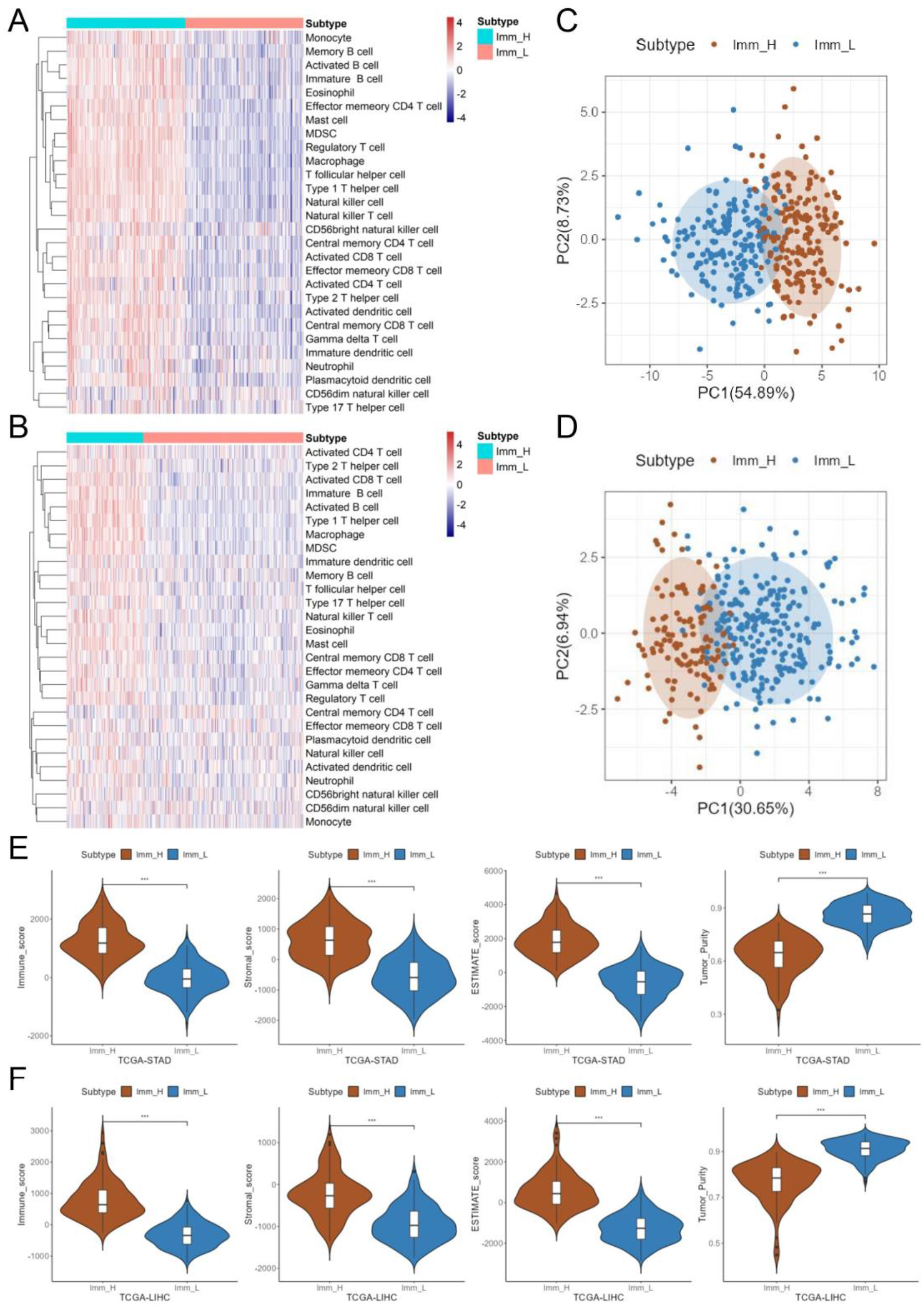
Identification and verification of immune subtypes in TCGA samples. (A-B) Heatmap of ssGSEA scores for 28 immune cell types of GC and HCC samples. Hierarchical clustering obtained two subtypes in TCGA-STAD and TCGA-LIHC datasets. (C-D) PCA of GC and HCC samples based on ssGSEA scores. Samples in different subtypes are represented by different colored dots. (E-F) Violin plot of tumor purity, ESTIMATE score, immune score, and stromal score of different subtypes in GC and HCC, respectively. \*\*\**P* < 0.001.

### 3.3 Immune characteristics of different immune subtypes

To further investigate the distinctions in immune cell types and proportions between the two subtypes described above, we estimated the proportion of 22 immune cell subsets in two immune subtypes utilizing the CIBERSORT algorithm. Results showed that for GC, the Imm_H group had significantly higher proportions of CD8+ T cell, M1 macrophage, M2 macrophage, and resting mast cell than the Imm_L group, but significantly lower proportions of M0 macrophage and activated mast cell than the Imm_L group (P<0.01) (Fig. 3A). As for HCC, the Imm_H group had significantly higher proportions of naive B cell, CD8+ T cell, M1 macrophage and lower proportions of resting NK cell and resting mast cell (P<0.01) (Fig. 3B). CD8+ T cells are widely recognized as a subset of immune cells that exert anti-tumor effect through cytotoxicity. However, recent findings have further subdivided CD8+ T cells into many subpopulations that secrete different cytokines and perform different functions [26]. The anti-tumor immune effect iduced by CD8+ T cells is also regulated by immune checkpoints, and activated immune checkpoint signaling can suppress the immune activity of CD8+ T cells. Immune checkpoints are often upregulated in the TME, leading to tumor immune escape [27]. Here we observed several immune checkpoint genes PDCD1 (PD-1), CD274 (PD-L1), PDCD1LG2, LAG3, CTLA4 and HAVCR2, and results showed that the expressions of all these immune checkpoints were significantly higher in the Imm_H group than that in the Imm_L group (P<0.01) (Fig. 3C-3H). These results indicated that although Imm_H group had more infiltrating immune cells, they were more likely to be in an immunosuppressive microenvironment, which prevent them from producing effective anti-tumor immune response.

**Fig. 3.**
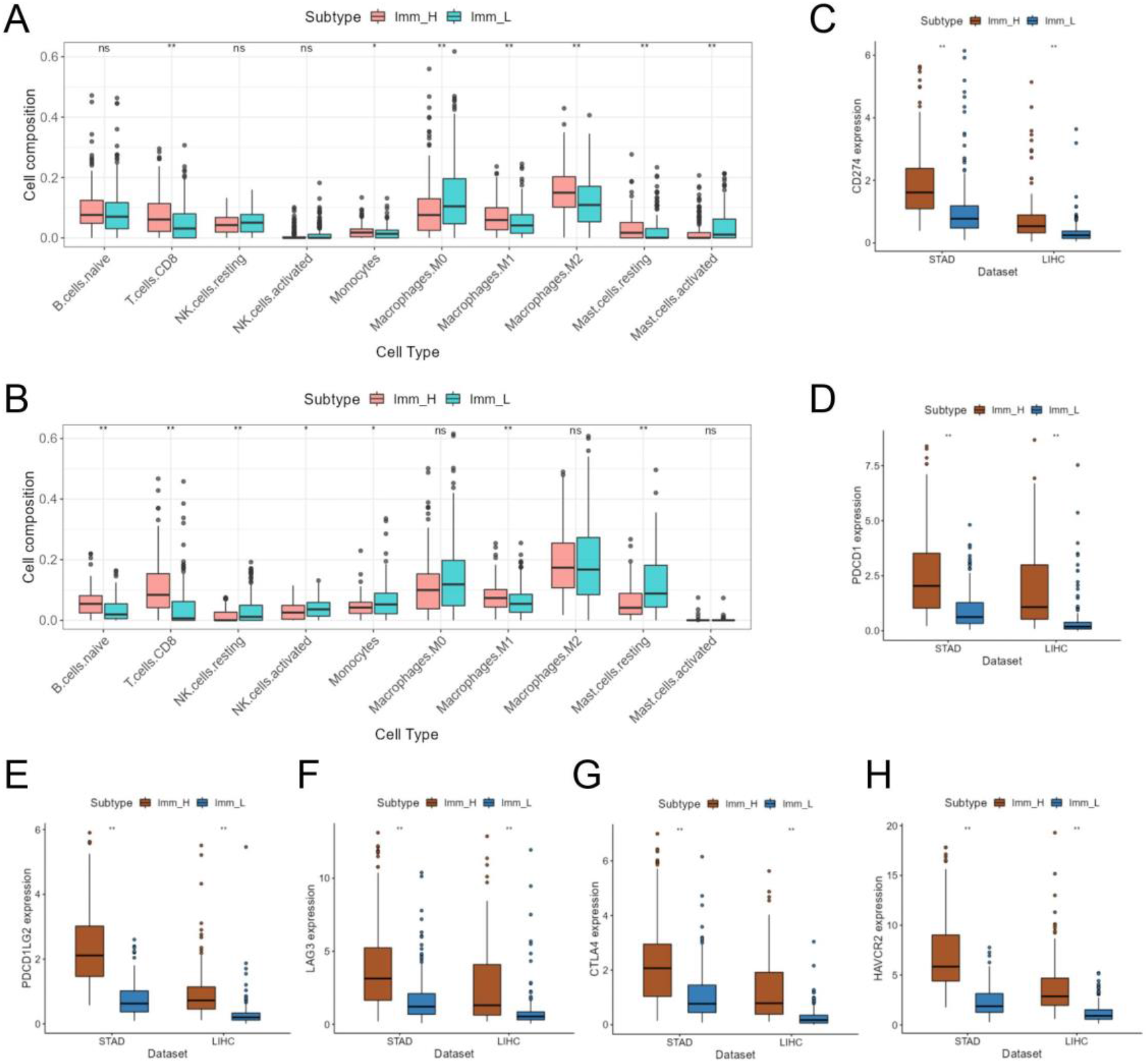
Different immunophenotypes of Imm_H and Imm_L subtypes. (A-B) Comparation of the proportion of immune cell subsets obtained by CIBERSORT between the Imm_H and Imm_L group in GC and HCC. (C-H) Expression of CD274, PDCD1, PDCD1LG2, LAG3, CTLA4, and HAVCR2 in different cancer and different subtypes. \**P*<0.05, \*\**P*<0.01.

### 3.4 Identification of differential genes, pathways and networks associated with immune subtypes

We identified genes differentially expressed between different immune subtypes in GC and HCC, respectively (Fig. 4A-4B). There were 2325 DEGs in TCGA-STAD and 1867 DEGs in TCGA-LIHC, and 914 shared DEGs between the two datasets (Fig. 4C). Next, common DEGs in GC and HCC were selected, and 196 differentially expressed immune genes (DEIGs) were screened out. DEIGs is obtained according to the gene list recorded in the ImmPort database (https://www.dev.immport.org), and each gene has its corresponding functional category. To further analyze the differential biological function and gene network among immune subtypes, we used OMIM database [28] and CIPHER method [29] to obtain GC and HCC related disease genes. The CIPHER algorithm was used to predict disease genes, and the top 1000 genes were considered as the prediction result. Genes that presented in both cancers were used as disease genes for subsequent analysis. After mapping disease genes and DEIGs to protein-protein interaction network from STRING (https://string-db.org/) [30], a group of genes closely related to disease genes were identified from these DEIGs, and protein interaction network among genes were constructed. There were 69 genes in the network, including disease genes as well as DEIGs overlapping in GC and HCC. We identify two main modules from the complete network (Fig. 4D-4E). Module 1 contains 24 genes, which are mainly associated with TCR signaling pathway, B cell receptor signaling pathway and natural killer cell cytotoxicity. Module 2 contains 27 genes, which are mainly related to cytokines and cytokines receptors.

**Fig. 4.**
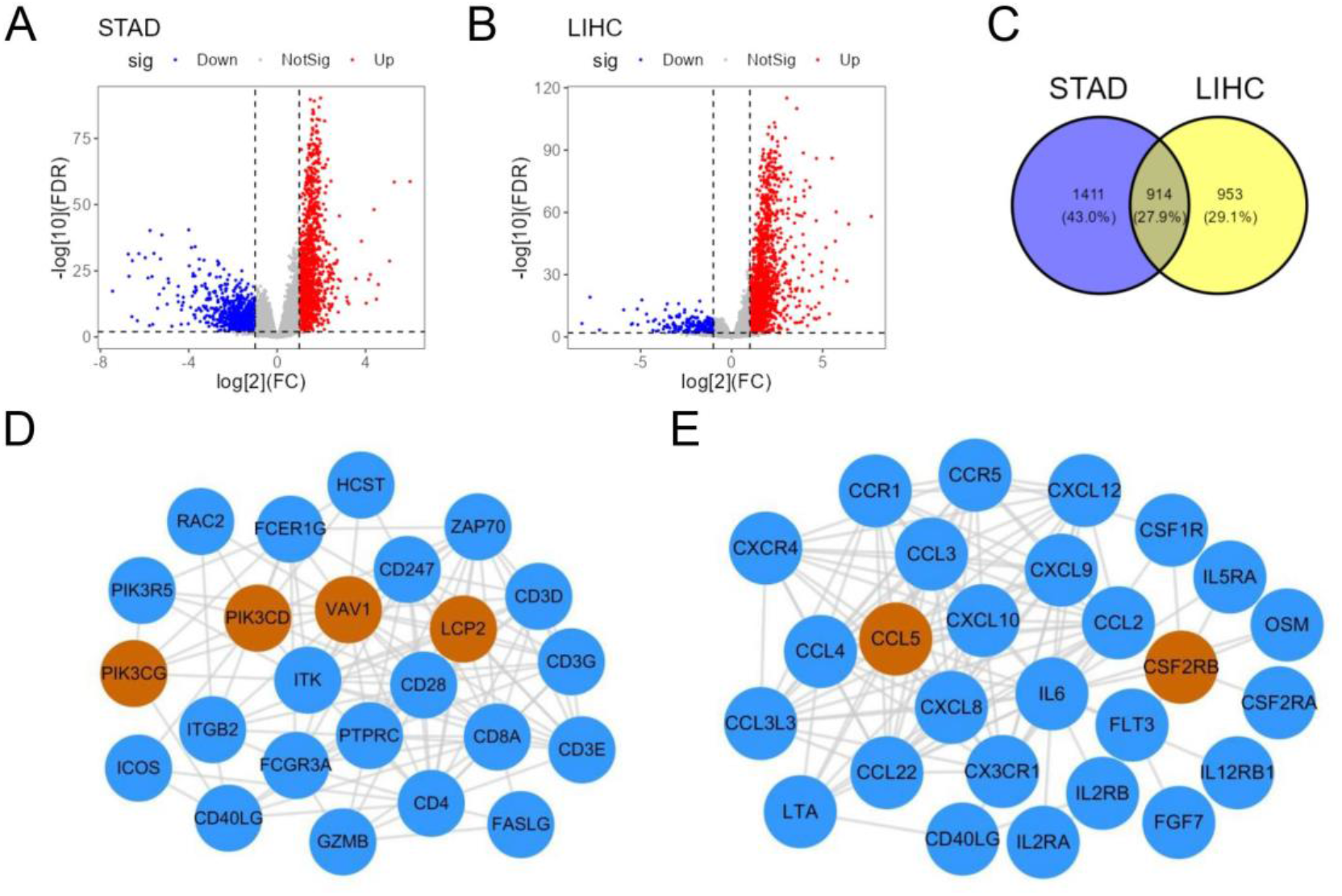
Differential genes networks associated with immune subtypes. Volcano plot shows the DEGs between Imm_H and Imm_L group in (A) TCGA-STAD and (B) TCGA-LIHC. (C) Venn diagram of DEGs between subtypes in TCGA-STAD and TCGA-LIHC. (D-E) Two key gene modules identified from the differential network. The blue and brown nodes represent the differentially expressed immune genes and disease genes obtained from OMIM and CIPHER, respectively.

### 3.5 Efficacy of synthetic OVs in mouse tumor models

We tested the inhibitory effect of synthetic oncolytic viruses expressing GM-CSF on tumor growth in two mouse tumor models. Similar to our previous results of OV-PD1 against Hepa1-6 tumor growth [19], significantly greater AGS tumor inhibition were observed for SynOV1.1 than the negative control (Fig.5A). IRTV of SynOV1.1 treated group reached 87%. No significant difference in bodyweight gain was observed between the control and SynOV1.1 treated group (data not shown). In the mHepa1-6 mouse syngeneic tumor model, SynOV1.1m appeared to inhibit tumor growth in a dose-dependent manner. Significantly greater tumor inhibition was observed for SynOV1.1m at the dose of 5 × 10^10^ VP than sorafenib (Fig.5B). IRTV reached 94% for SynOV1.1m at the dose of 5×10^10^ VP as compared to 68% for sorafenib at the dose of 60 mg/kg. No significant difference in bodyweight gain was observed between the control and SynOV1.1m treatment groups (data not shown).

**Fig. 5.**
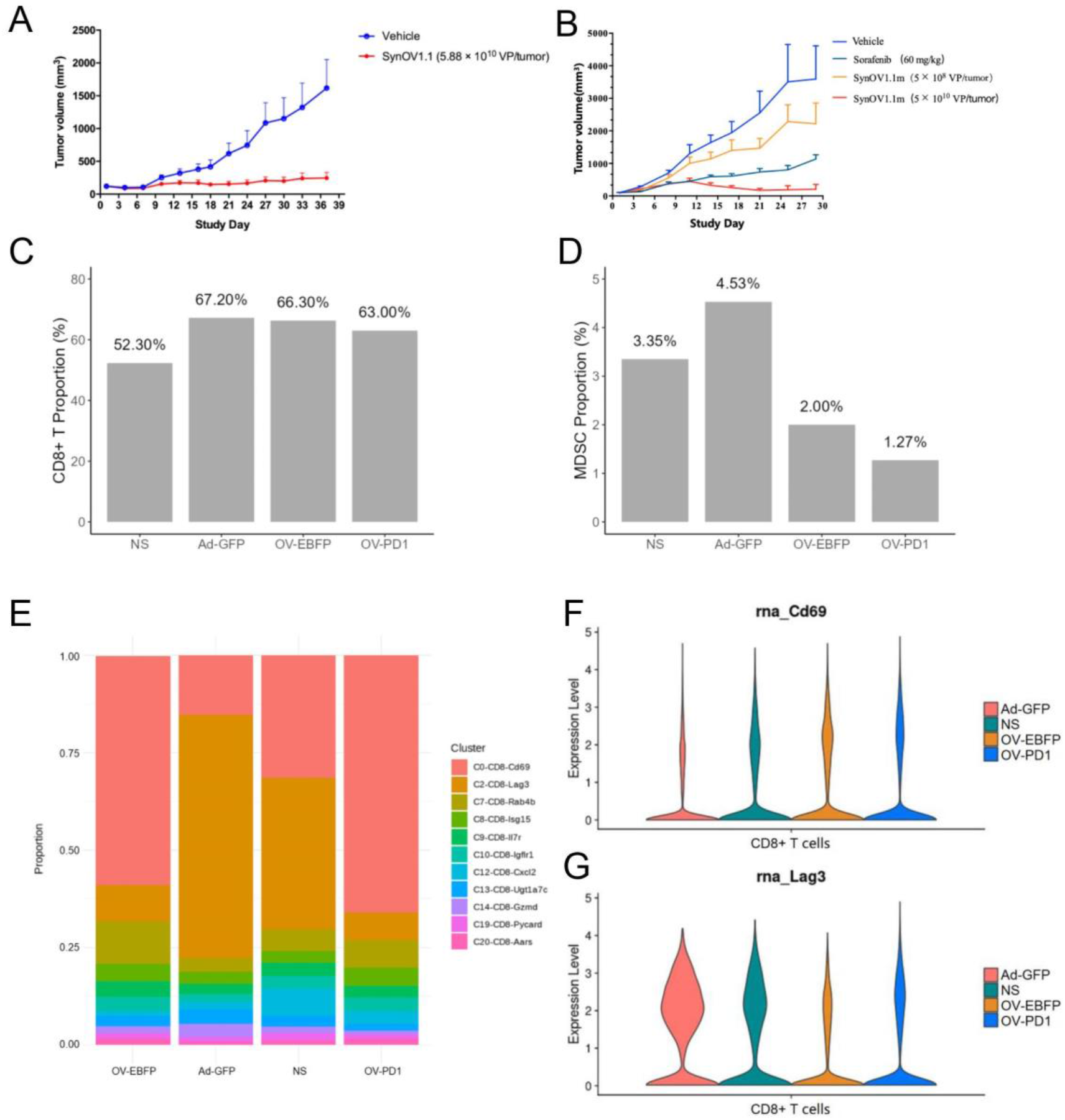
Effect of synthetic oncolytic virus on tumor growth and tumor microenvironment in mice. (A) Inhibition of AGS tumor growth by SynOV1.1 intratumoral injection. (B) Inhibition of mHepa1-6 tumor growth by SynOV1.1m intratumoral injection. (C-D) Proportion of CD8+ T cells and MDSCs in tumors of different groups. (E) Proportion of CD8+ T cell subpopulations in different groups. (F-G) Expression of Cd69 and Lag3 mRNA in CD8+ T cells in different groups.

To investigate the effect of synthetic oncolytic virotherapy on tumor microenvironment, we established hepa1-6 tumor model in immune-competent mice. On the 10th day after treatment with AD-GFP, OV-EBFP, OV-PD1 and normal saline (NS group), mouse tumors were dissected for single-cell RNA sequencing. After clustering and cell type identification of the single-cell data, it was found that the proportion of CD8+ T cells in the three experimental groups after virus injection was significantly higher than that in the control group, which reflected the promoting effect of oncolytic virus on CD8+ T cell infiltration (Fig. 5C). CD11b+GR-1+ myeloid-derived suppressor cells (MDSCs) in different experimental groups were identified and the proportion of MDSCs in total cells was calculated. It was found that the proportion of MDSCs decreased significantly after oncolytic virus treatment, and the effect of synthetic oncolytic virus expressing immune checkpoint inhibitor on MDSC was more obvious (Fig. 5D).

Although injection of all three viruses promoted infiltration of CD8+ T cells, the type and status of CD8+ T cells may differ and require further analysis. We further calculated the proportion of CD8+ T cell subsets in different groups. CD8+ T cells expressing Lag3 were predominant in NS group and AD-GFP group, while CD8+ T cells expressing Cd69 were predominant in OV-EBFP and OV-PD1 group (Fig. 5E). Cd69 is an important marker of early activation of effector T cells. LAG3 is a membrane protein with immunosuppressive function, which can bind to type II MHC molecules expressed by tumor cells and prevent the normal TCR recognition mechanism. Oncolytic virus treatment increased Cd69 expression and decreased Lag3 expression in CD8+ T cells (Fig.5F-5G). After oncolytic virus treatment, the proportion of effector CD8+ T cell subtypes increased while the proportion of immunosuppressive T cell subtypes decreased, and the effect of synthetic oncolytic virus expressing immune checkpoint inhibitors was more prominent. However, the proportion of immunosuppressive T cell subtypes in the tumor after injection of the non-replicating adenovirus (Ad-GFP) increased in a certain extent compared to the NS group, which promoted immunosuppression in the tumor immune microenvironment.

## 4. Discussion

Gastric cancer and hepatocellular carcinoma are both common cancers, and their clinical needs are currently unmet. Despite the fact that cancer immunotherapy has succeeded in certain cancers, only a small percentage of GC and HCC patients have satisfying response to immunotherapy. Many studies have proposed that the response to immunotherapy and tumor immune microenvironment are strongly connected [12,31]. Our research showed that the type and proportion of tumor-infiltrating immune cells differed notably among different samples, and the samples could be clustered into two subtypes according to the immune infiltration score. In addition, the estimated tumor purity was significantly different between the two subtypes with different immune infiltration level. The infiltration of almost all immune cell subtypes were higher in the Imm_H group, including anti-tumor and pro-tumor /immunosuppressive immune cells. The Imm_H group also had higher expression of CD274, PDCD1 and several other immune checkpoint genes. As CD8+ T cells infiltration and PD-L1 expression are positively correlated to patients’ outcome after immunotherapy [32,33], the results suggest that patients in Imm_H group are more likely to have satisfying response to PD-1/PD-L1 targeting therapy.

Previous experiments demonstrated that the synthetic oncolytic viruses OV-EBFP and OV-PD1 suppressed tumor growth and increased the fraction of IFN-γ+ and Ki-67+ cells in tumor infiltrating CD8+ T cells in immune-competent mouse tumor model [19]. In this work, we used single-cell RNA sequencing to further assess the type and proportion of immune cells in the TME. The proportion of CD8+ T cells notably increased after OV-PD1 treatment while the proportion of MDSCs significantly reduced. The beneficial anti-tumor immune responses induced by T cells and NK cells can be strongly inhibited by MDSCs, which can accelerate tumor growth and weaken the effect of immunotherapy. Growth factors secreted by tumors can recruit MDSCs to tumor tissues, and these MDSCs promote tumorigenesis through kinds of mechanisms inducting T cell and NK cell exhaustion [34]. A meta-analysis found that MDSC enrichment in solid tumor was connected to poor prognosis and overall survival [35]. OV-PD1 treatment also altered the proportion of different subsets of CD8+ T cells. These findings indicated that the mechanism of OV-PD1 in different immune subtypes may be different. It may decrease the proportion of immunosuppressive cells and inhibit PD-1/PD-L1 pathway in the Imm_H group while promoting CD8+ T cells infiltration in the Imm_L group. The tumor microenvironment contains various kinds of immune cells in addition to CD8+ T cells and MDSCs, and the effect of oncolytic virus on other immune cells needs to be further studied.

Traditional Chinese medicine (TCM) has been used in treating cancer for a long time and has accumulated a large amount of clinical experience [36]. The important theory of TCM syndrome (also known as ZHENG) embodies the unique holistic view of TCM [37], and TCM plays an important role in the treatment of many complex diseases [38]. The use of TCM in cancer is becoming acknowledged all over the world as more and more evidences of biological mechanism and clinical trials are accumulated [39,40]. Health-strengthening (*Fu-Zheng*) herbs have been utilized widely in cancer treatment and many researches have been conducted on the mechanism of health-strengthening herbs from the perspectives of immunity and metabolism [41,42]. A previous research combining network pharmacology and high-throughput experiments showed that health-strengthening herbs are more inclined to regulate tumor immunity than directly kill tumor cells, which reflects the great potential of health-strengthening herbs in anti-tumor immunity [43]. Further research on the effects of health-strengthening TCM on different immune subtypes of cancer will help us to further understand their biological mechanism.

In conclusion, our study divides GC and HCC into two subtypes with different immune infiltration. The effect of synthetic oncolytic adenovirus on tumor growth and tumor immune microenvironment was explored in mouse models, and the potential mechanism of OV-PD1 treatment for both subtypes was analyzed.

